# The FlhA linker mediates flagellar protein export switching during flagellar assembly

**DOI:** 10.1101/2020.09.09.290478

**Authors:** Yumi Inoue, Mamoru Kida, Miki Kinoshita, Norihiro Takekawa, Keiichi Namba, Katsumi Imada, Tohru Minamino

**Author notes:** Department of Ophthalmology and Visual Sciences, Kyoto University Graduate School of Medicine, Kyoto, 606-8507 Japan (Yumi Inoue). Correspondence to Tohru Minamino : Mailing address, Graduate School of Frontier Biosciences, Osaka University, 1-3 Yamadaoka, Suita, Osaka 565-0871, Japan.; or Katsumi Imada: Graduate School of Sciences, Osaka University, 1-1 Machikaneyama, Toyonaka, Osaka 560-0043, Japan.

## Abstract

The flagellar protein export apparatus switches export specificity from hook-type to filament-type upon completion of hook assembly, thereby initiating filament assembly at the hook tip. The C-terminal cytoplasmic domain of FlhA (FlhA_C_) forms a homo-nonameric ring structure that serves as a docking platform for flagellar export chaperones in complex with their cognate filament-type substrates. Interactions of the flexible linker of FlhA (FlhA_L_) with its nearest FlhA_C_ subunit in the ring allow the chaperones to bind to FlhA_C_ to facilitate filament-type protein export, but it remains unclear how it occurs. Here, we report that FlhA_L_ acts as a switch that brings the order to flagellar assembly. The crystal structure of FlhA_C_(E351A/D356A) showed that Trp-354 in FlhA_L_ bound to the chaperone-binding site of its neighboring subunit. We propose that FlhA_L_ binds to the chaperon-binding site of FlhA_C_ to suppress the interaction between FlhA_C_ and the chaperones until hook assembly is completed.

## Introduction

The flagellum of *Salmonella enterica* (hereafter referred to *Salmonella*) is a supramolecular motility machine consisting of the basal body, the hook and the filament. For construction of the flagella on the cell surface, a type III protein export apparatus (fT3SS) transports flagellar building blocks from the cytoplasm to the distal end of the growing flagellar structure. The fT3SS is divided into three structural parts: a transmembrane export gate complex made of FlhA, FlhB, FliP, FliQ and FliR, a docking platform composed of the cytoplasmic domains of FlhA and FlhB (FlhA_C_ and FlhB_C_), and a cytoplasmic ATPase ring complex consisting of FliH, FliI and FliJ
^1^. The FlhA_C_-FlhB_C_ docking platform switches its substrate specificity from hook-type export substrates (FlgD, FlgE and FliK) to filament-type ones (FlgK, FlgL, FlgM, FliC and FliD) when the hook reaches its mature length of about 55 nm in *Salmonella*, thereby terminating hook assembly and initiating filament formation^2^.

FliK is secreted via fT3SS during hook-basal body (HBB) assembly not only to measure the hook length but also to switch export specificity of the FlhA_C_-FlhB_C_ docking platform^2^. This has been recently verified by *in vitro* reconstitution experiments using inverted membrane vesicles^3,4^. The N-terminal domain of FliK (FIiK_N_) serves as a secreted molecular ruler to measure the hook length^5–8^. When the hook length reaches about 55 nm, a flexible linker region of FliK connecting FIiK_N_ and the C-terminal domain (FliK_C_) promotes a conformational rearrangement of FliK_C_ to interact with FlhB_C_, thereby terminating hook-type protein export^9,10^.

FlhA_C_ consists of four domains, D1, D2, D3 and D4, and a flexible linker (FlhA_L_) connecting FlhA_C_ with the N-terminal transmembrane domain of Flh_A_ (Fig. 1)^11^. FlhA_C_ forms a homo-nonamer ring^12,13^ and provides binding-sites for flagellar export chaperons (FlgN, FliS and FliT) in complex with their cognate filament-type substrates^14–17^. Interactions of FlhA_C_ with the flagellar chaperones facilitate the filament-type substrates to enter into the export gate complex for efficient protein export and assembly^15,16^. The FliK_C_-FlhB_C_ interaction is thought to induce structural remodeling of the FlhA_C_ ring through interactions of FlhA_L_ with its nearest FlhA_C_ subunit, thereby initiating the export of filament-type proteins^13,18,19^. However, it remains unknown how.

**Fig. 1.**
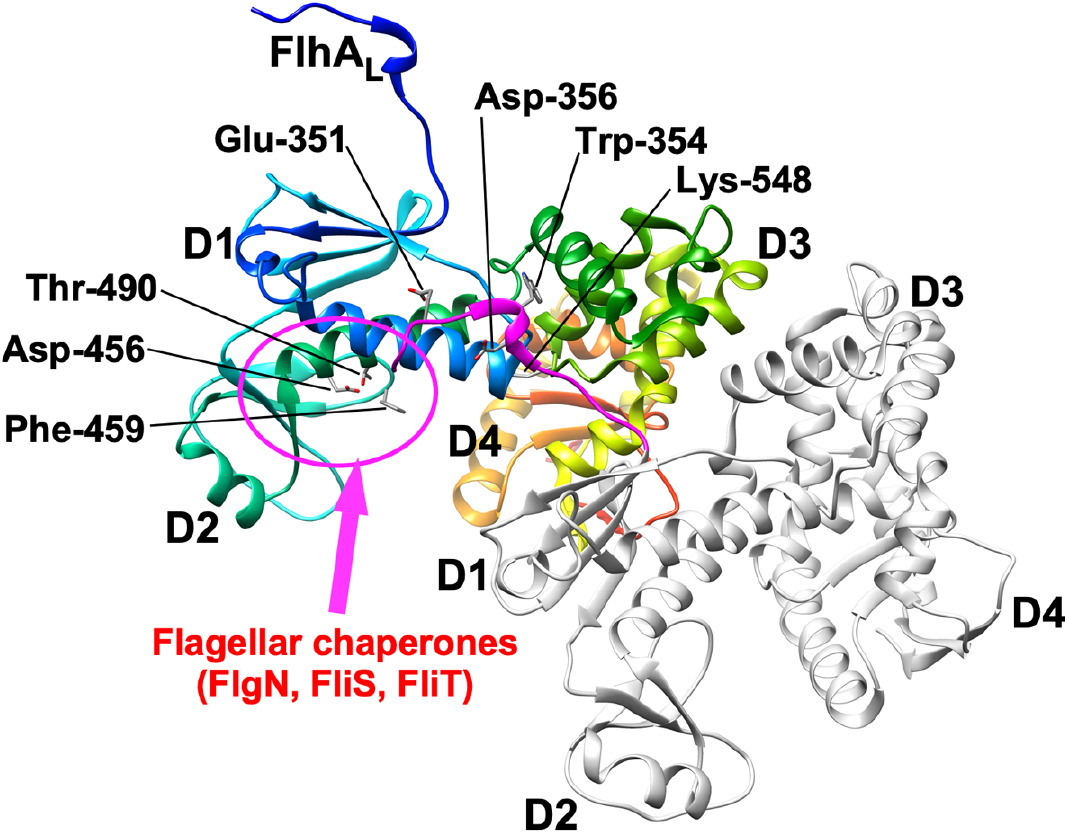
FlhA_C_ structure in a crystal (PDB ID: 3A5I). FlhA_C_ consists of four domains, D1, D2, D3 and D4 and a flexible linker (FlhA_L_). Glu-351, Trp-354 and Asp-356 of FlhA_L_ binds to the D1 and D3 domains of its neighboring subunit. A well-conserved hydrophobic dimple including Phe-459 is responsible for the interaction of FlhA_C_ with flagellar export chaperones in complex with filament-type substrates. Phe-459 and Lys-548 are exposed to solvent on the molecular surface when FlhA_C_ adopts the open conformation. The interactions between the two FlhA_C_ molecules in this crystal also represent those in the FlhA_C_ nonameric ring of the export apparatus.

In the present study, to clarify the role of FlhA_L_ in the export switching mechanism of fT3SS, we analyzed the interaction between FlhA_L_ and FlhA_C_ and provide evidence suggesting that the interaction of FlhA_L_ with the chaperone-binding site of FlhA_C_ inhibits the binding of the flagellar export chaperones to FlhA_C_ to keep the export specificity in the hook type during hook assembly.

## Results

### Isolation of pseudorevertants from the *flhA(E351A/W354A/D356A)* mutant

Glu-351, Trp-354 and Asp-356 of FlhA_L_ bind to the D1 and D3 domains of its neighboring FlhA_C_ subunit to stabilize FlhA_C_ ring structure in solution^11,13^. The *flhA(E351A/D356A)* and *flhA(W354A)* mutants produces the HBBs without the filament attached although their hook length is not controlled properly^13^. The W354A and E351A/D356A mutations inhibit the interaction of FlhA_C_ with flagellar chaperones in complex with their cognate filament-type substrates, suggesting that the interaction between FlhA_L_ and the D1 and D3 domains of its neighboring FlhA_C_ subunit keeps the chaperone binding site of FlhA_C_ open to allow the chaperones to bind to FlhA_C_ to facilitate the export of the filament-type substrates^13^. However, the *flhA(E351A/W354A/D356A)* mutant do not produce the HBBs at all, raising a question of why the E351A/W354A/D356A triple mutation inhibits HBB assembly^13^. To clarify this question, we isolated 14 pseudorevertants from the *flhA(E351A/W354A/D356A)* mutant. Motility was somewhat restored by these pseudorevertant mutations but it was much poorer than that of wild-type cells (Fig. 2a). Export substrates such as FlgD, FlgE, FlgK and FliD were detected in the culture supernatants of these pseudorevertants (Fig. 2b). In agreement with this, these pseudorevertants produced a couple of flagella on the cell surface (Fig. 2c). DNA sequencing revealed that all suppressor mutations are located in the *flgMN* operon. One was the M1I mutation at the start codon of the *flgM* gene (isolated twice), presumably inhibiting FlgM translation. Two suppressor mutations produced a stop codon at position of Gln-52 or Ser-85 of FlgM, resulting in truncation of the C-terminal region of FlgM. Nine suppressor mutations were large deletions in *flgM.* We also found that there was a large deletion in the *flgM* and *flgN* genes, thereby disrupting both FlgM and FlgN. A loss-of-function of FlgM results in a considerable increment in the transcription levels of flagellar genes^20,21^. Consistently, the cellular levels of FlgK and FliD seen in the pseudorevertants were higher than those in its parental strain (Fig. 2b, 3rd and 4th rows). It has been shown that an interaction between FliJ and FlhA_L_ brought about by FliH and FliI fully activates the transmembrane export gate complex of fT3SS to utilize proton motive force (PMF) across the cell membrane to drive flagellar protein export^22^. Because the E351A/W354A/D356A triple mutation reduces the binding affinity of FlhA_C_ for FliJ^13^, this suggests that these *flgM* mutations considerably increase the cytoplasmic levels of FliH, FliI, FliJ and export substrates to allow the *flhA(E351A/W354A/D356A)* mutant to export flagellar building blocks for producing a small number of flagella on the cell surface. Therefore, we propose that Glu-351, Trp-354 and Asp-356 of FlhA_L_ also play an important role in the activation mechanism of the PMF-driven export gate complex.

**Fig. 2.**
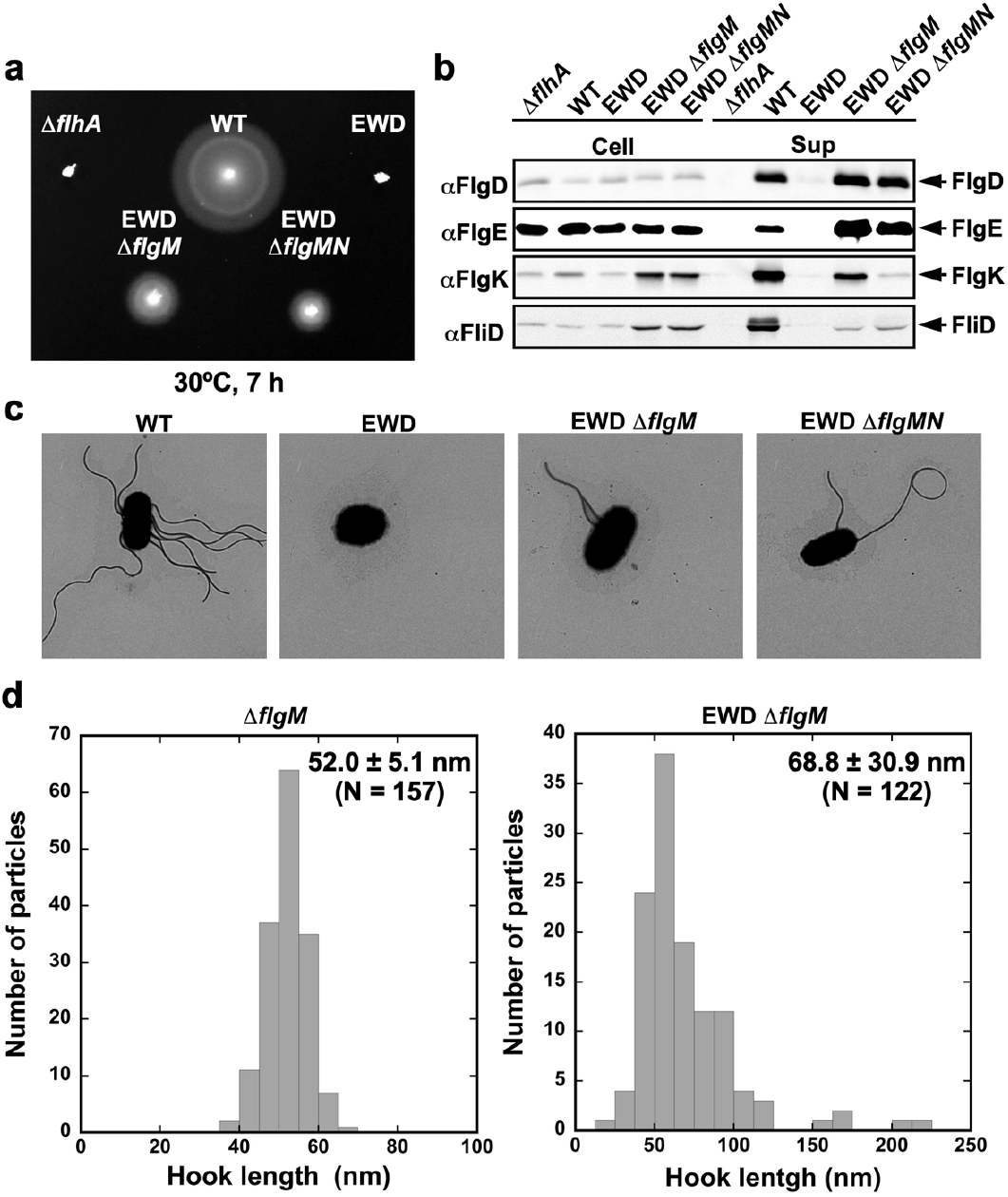
Isolation pseudorevertants from the *flhA(E351A/W354A/D356A)* mutant. (a) Motility of the *Salmonella* NH001 strain transformed with pTrc99A *(ΔflhA)*, pMM130 (WT), or pYI003 [FlhA(E351A/W354A/D356A) indicated as EWD]), YI1003-4 (EWD Δ*flgM*) or YI1003-13 (EWD *ΔflgMN)* in soft agar. (b) Immunoblotting using polyclonal anti-FlgD (1st row), anti-FlgE (2nd row), anti-FlgK (3rd row) or anti-FliD (4th row) antibody, of whole cell proteins (Cell) and culture supernatants (Sup) prepared from the above strains. (c) Electron micrographs of the above cells. (d) Histogram of hook length distribution of NH001gM (Δ*flhA ΔflgM::km)* carrying pMM130 (Δ*flgM*) or pYI003 (EWD *ΔflgM).*

We found that the secretion levels of FlgD and FlgE by the pseudorevertants were about 1.5-fold and 4-fold higher than those by the wild-type whereas the secretion levels of FlgK and FliD were much lower (Fig. 2b), raising the possibility that *flgM* suppressor mutations do not efficiently promote export switching of fT3SS from hook-type substrates to filament-type ones in the *flhA(E351A/W354A/D356A)* mutant. To clarify this, we introduced the *DflgM::km* allele to the *Salmonella* NH001 *(DflhA)* strain to produce the *DflgM::km* and *flhA(E351A/W354A/D356A) DflgM::km* cells (Supplementary Fig. 1). The *DflgM::km* allele restored motility of the *flhA(E351A/W354A/D356A)* mutant in a way similar to other *flgM* suppressor mutations. Then, we isolated flagella from the *DflgM::km* and *flhA(E351A/W354A/D356A) DflgM::km* cells and measured their hook length. The hook length of the *DflgM::km* strain was 52.0 ± 5.1 nm (mean ± SD, n = 157) (Fig. 2d), which is nearly the same as that of the wild-type strain (51.0 ± 6.9 nm)^13^. This indicates that the loss-of-function mutation of FlgM does not affect the hook length control. In contrast, the average hook length of the *flhA(E351A/W354A/D356A) DflgM::km* strain was 68.8 ± 30.9 nm (mean ± SD, n = 157) (Fig. 2d), indicating that the hook length control becomes worse in the presence of the E351A/W354A/D356A triple mutation. These suggest that this triple mutation affects not only the initiation of filament-type protein export but also the termination of hook-type protein export. Therefore, we propose that conformational rearrangements of FlhA_L_ are required for well-regulated export switching of fT3SS.

### Effect of FlhA linker mutations on the hydrodynamic properties of FlhA_C_ in solution

A well conserved hydrophobic dimple of FlhA_C_ containing Asp-456, Phe-459 and Thr-490 resides is located at the interface between domains D1 and D2 and is involved in the interactions with the FlgN, FliS and FliT chaperones in complex with their cognate filament-type substrates (Fig. 1)^15–17^. The W354A, E351A/D356A and E351A/W354A/D356A mutations significantly reduce the binding affinity of FlhA_C_ for these chaperone/substrate complexes^13^, raising the possibility that FlhA_L_ carrying these *flhA* mutations binds to the hydrophobic dimple of FlhA_C_ and blocks the FlhA_C_-chaperone interaction. If true, FlhA_C_ with these mutations would show distinct hydrodynamic properties compared with wild-type FlhA_C_. To clarify this possibility, we performed size exclusion chromatography with a Superdex 75 column HR 10/30 column. Wild-type His-FlhA_C_ appeared as a single peak at an elution volume of 10.3 mL, which corresponds to the deduced molecular mass of His-FlhA_C_ (about 43 kDa) (Fig. 3a). His-FlhA_C_(W354A), His-FlhA_C_(E351A/D356A) and His-FlhA_C_(E351A/W354A/D356A) appeared as a single peak at an elution volume of 10.3 mL, 10.5 mL and 10.4 mL, respectively (Fig. 3a), indicating that these mutant variants exist as a monomer in solution. His-FlhA_C_(E351A/D356A) exhibited a slightly delayed elution behavior compared with the wild-type. Furthermore, His-FlhA_C_(E351A/D356A) showed a slightly faster mobility on SDS-PAGE gels. Far-UV CD measurements revealed that the E351A/D356A double mutation did not affect the secondary structures of FlhA_C_ (Supplementary Fig. 2). These suggest that FlhA_C_(E351A/D356A) adopts a more compact conformation than wild-type FlhA_C_. The elution peak position of His-FlhA_C_(E351A/W354A/D356A) was between those of the wild-type and FlhA_C_(E351A/D356A) (Fig. 3a). Because His-FlhA_C_(E351A/W354A/D356A) showed two different bands on SDS-PAGE gels, with a slower mobility band corresponding to wild-type FlhA_C_ and a faster one corresponding to FlhA_C_(E351A/D356A) (Fig. 3a, inset), we suggest that FlhA_C_(E351A/W354A/D356A) exists in an equilibrium between the wild-type conformation and the compact conformation. Since FlhA_C_(W354A) adopted the wild-type conformation (Fig. 3a), we suggest that the E351A/D356A double mutation is required to make FlhA_C_ more compact and that Trp-354 of FlhA_L_ is needed to stabilize the compact conformation.

**Fig. 3.**
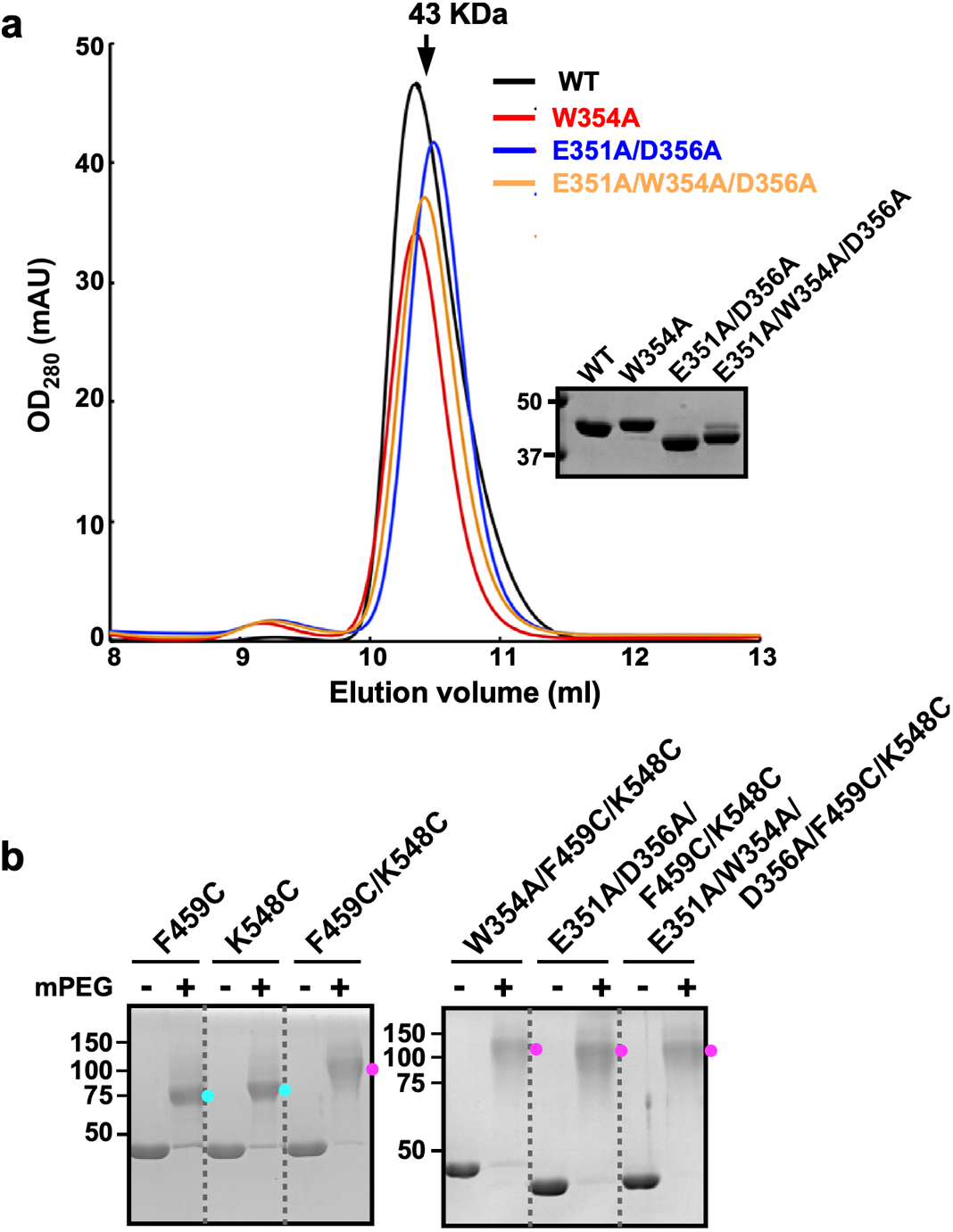
Effect of FlhA linker mutations on the FlhA_C_ conformation. (a) Size exclusion chromatography using a Superdex 75HR 10/30 column. Elution positions of His-FlhA_C_ (WT, black), His-FlhA_C_(W354A) (red), His-FlhA_C_(E351A/D356A) (blue) and His-FlhA_C_(E351A/W354A/D356A) (orange) are 10.3 ml, 10.3 ml, 10.5 ml and 10.4 ml, respectively. Arrow indicates the elution peak of ovalbumin (43 kDa). Inset, CBB-stained gels of purified FlhA_C_ proteins. (b) Effect of FlhA mutations on mPEG-maleimide modification of Cys459 and Cys548. His-FlhA_C_(F459C), His-FlhA_C_(K548C), His-FlhA_C_(F459C/K548C), His-FlhA_C_(W354A/F459C/K548C), His-FlhA_C_(E351A/D356A/F459C/K548C) and His-FlhA_C_(E351A/W354A/D356A/F459C/K548C) were incubated with (+) or without (-) mPEG-maleimide. After centrifugation at 20,000 g for 20 min to remove any aggregates, supernatants were analyzed by SDS-PAGE with CBB staining. Cyan and magenta dots indicate positions of FlhA_C_-(mPEG) and FlhA_C_-(mPEG)_2_, respectively.

### Effect of FlhA linker mutations on methoxypolyethylene glycol 5000 maleimide (mPEG-maleimide) modifications of Cys-459 and Cys-548

FlhA_C_ structures have shown that it adopts three distinct, open, semi-closed and closed conformations^11,14,17,18,23^. Phe-459 and Lys-548 are fully exposed to solvent on the molecular surface of the open conformation of FlhA_C_ but are in close proximity to each other in the closed conformation^11,18,23^. To test whether mutations in FlhA_L_ bias FlhA_C_ towards the closed structure, we performed Cys modification experiments with mPEG-maleimide, which adds a molecular mass of ~5 kDa to a target protein. FlhA_C_(F459C/K548C) modified by mPEG-maleimide showed much slower mobility shift, indicating that both Cys459 and Cys548 are exposed to the solvent. The W354A, E351A/D356A and E351A/W354A/D356A mutations did not inhibit Cys modifications with mPEG-maleimide at all, indicating that FlhA_C_ with these mutations does not adopt the closed conformation.

### Crystal structure of FlhA_C_(E351A/D356A)

To investigate whether Trp-354 of FlhA_L_ binds to the hydrophobic dimple of FlhA_C_ to makes FlhA_C_(E351A/D356A) more compact, we explored crystallization conditions of FlhA_C_(E351A/D356A) for a molecular packing distinct from the open (PDB code: 3A5I)^11^ and semi-closed (PDB code: 6AI0)^18^ forms of wild-type FlhA_C_. We found a new orthorhombic crystal that diffracted up to 2.8 Å resolution, with unit cell dimensions *a* = 71.7 Å, *b* = 96.2 Å, *c* = 114.1 Å (Table 1) and the asymmetric unit containing two FlhA_C_ molecules (A and B). Mol-A adopts an open conformation similar to the 3A5I structure whereas Mol-B shows a semi-closed conformation similar to the 6AI0 structure (Supplementary Fig. 3). The residues from Val-349 to Val-357 in FlhA_L_ of Mol-A form an α-helix, which interacts with the hydrophobic dimple of a neighboring Mol-A molecule related by a crystallographic symmetry (Fig. 4a). Trp-354 fits into the hydrophobic dimple, and Ala-351 hydrophobically contacts with Pro-442 on the periphery of the dimple (Fig. 4b). These interactions resemble the interaction between the N-terminal α-helix of FliS and the hydrophobic dimple of FlhA_C_ (PDB ID: 6CH3)^17^ (Fig. 4c). Ile-7 and Tyr-10 of the N-terminal α-helix of FliS is in the corresponding position of Ala-351 and Trp-354 of FlhA_L_, respectively. Tyr-10 fits into the hydrophobic dimple of FlhA_C_, and Ile-7 interacts with Pro-442 of FlhA_C_ (Fig. 4d). These observations suggest that FlhA_L_ and flagellar chaperones bind competitively to a common binding site on FlhA_C_ and that the dissociation of FlhA_L_ from this binding site is required for the binding of the flagellar chaperones to FlhA_C_.

**Fig. 4.**
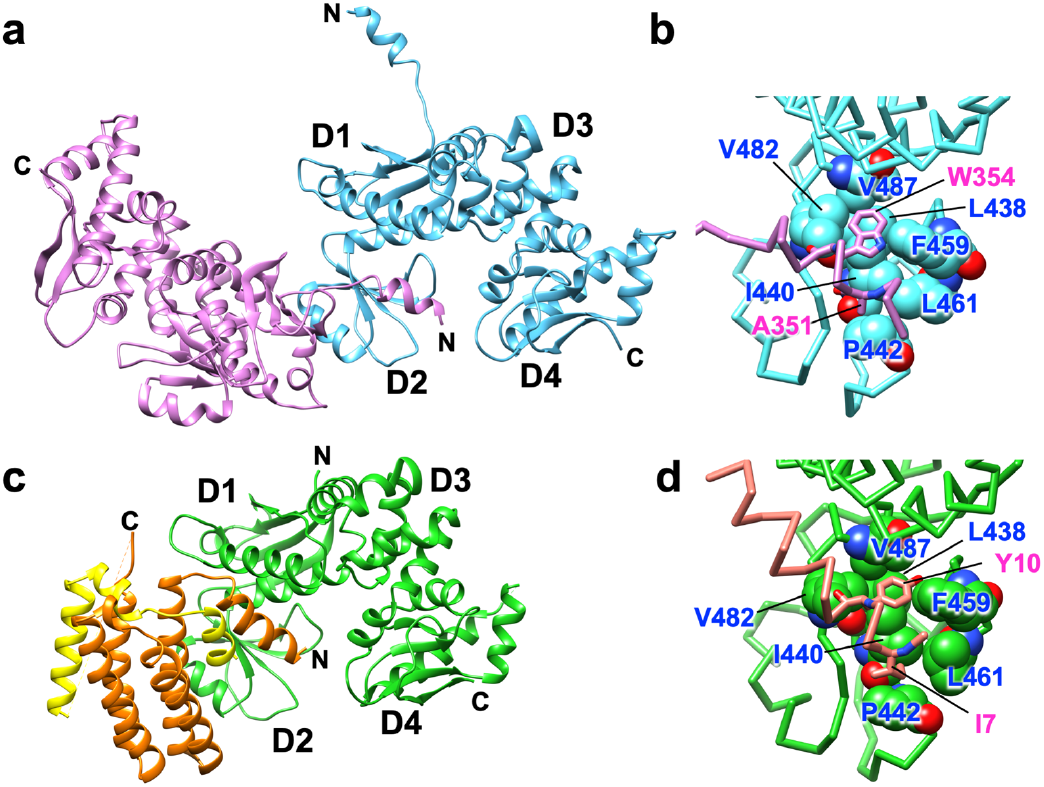
Interaction between FlhA_L_ and a well conserved hydrophobic dimple of its neighboring FlhA_C_ in the crystal of FlhA_C_(E351A/D356A). (a) Mol-A of FlhA_C_(E351A/D356A) (magenta) interacts with neighboring Mol-A (cyan) related by a crystallographic symmetry. (b) Close-up view of the interaction between FlhA_L_ and the hydrophobic dimple. Residues that form the hydrophobic dimple are indicated by balls. Ala-351 and Trp-354 in FlhA_L_ are shown in stick models. (c) Interaction between FlhA_C_ (green) and FliS (orange) fused with the C-terminal region of FliC (yellow) (PDB code: 6CH3). (d) Close-up view of the interaction between FliS and the hydrophobic dimple. The residues that form the hydrophobic dimple are indicated by ball. Ile-7 and Tyr-10 of FliS are shown in stick models.

**Table 1.** X-ray data collection and refinement statistics

| **X-ray data collection** |  |  |
| --- | --- | --- |
| Space group | *P2<sub>1</sub>2<sub>1</sub>2<sub>1</sub>* |  |
| Cell dimensions<br>*a*, *b*, *c* (Å) | 71.7, 96.2, 114.1 |  |
| Wavelength (Å) | 1.0000 |  |
| Resolution (Å) | 73.5-2.80 | (2.95-2.80) |
| *R*<sub>merge</sub> | 0.074 | (0.317) |
| *R*<sub>pim</sub> | 0.065 | (0.283) |
| CC(1/2) | 0.995 | 0.915 |
| *I*/σ*I* | 8.1 | (2.8) |
| Completeness (%) | 97.1 | (94.6) |
| Redundancy | 3.4 | (3.1) |
| **Refinement statistics** |  |  |
| Resolution range (Å) | 73.5-2.80 | (2.87-2.80) |
| No. of reflections working | 17,398 | (1,182) |
| No. of reflections test | 1,903 | (112) |
| *R*<sub>w</sub> (%) | 23.2 | (33.5) |
| *R*<sub>free</sub> (%) | 29.0 | (42.1) |
| Rms deviation bond length (Å) | 0.003 |  |
| Rms deviation Bond angle (°) | 0.680 |  |
| B-factors |  |  |
| Protein atoms | 70.0 |  |
| Solvent atoms | - |  |
| Ramachandran plot (%) |  |  |
| Most favored | 96.0 |  |
| Allowed | 3.9 |  |
| Disallowed | 0.1 |  |
| No. of protein atoms | 5, 252 |  |
| No. of solvent atoms | 0 |  |

Values in parentheses are for the highest resolution shell.
$R_w = \sum ||F_o| - |F_c|| / \sum |F_o|$ , $R_{free} = \sum ||F_o| - |F_c|| / \sum |F_o|$

## Discussion

The FlhA_C_ ring serves as the docking platform for flagellar export chaperones in complex with their cognate substrates and facilitates the export of filament-type proteins to form the filament at the hook tip after completion of hook assembly^14–17^. The FlhA_C_ ring also ensures the strict order of flagellar protein export, thereby allowing the huge and complex flagellar structure to be built efficiently on the cell surface^13,14,16,18,19^. Interactions of FlhA_L_ with its neighboring FlhA_C_ subunit in the nonamer ring is required for the initiation of filament-type protein export upon completion of hook assembly. However, it remained unclear how the FlhA_C_ ring mediates such hierarchical protein export during flagellar assembly.

In this study, we first performed genetic analyses of the *flhA(E351A/W354A/D356A)* mutant and found that the E351A/W354A/D356A triple mutation caused a loose hook-length control (Fig. 2d), indicating that the E351A/W354A/D356A mutation significantly affects the termination of hook-type protein export. Furthermore, this triple mutation also reduced the secretion levels of the filament-type proteins considerably (Fig. 2b), thereby reducing the number of flagellar filaments per cell (Fig. 2c). These results suggest that FlhA_L_ serves as a structural switch for substrate specificity switching of fT3SS from hook-type to filament-type and that Glu-351, Trp-354 and Asp-356 of FlhA_L_ are directly involved in this export switching mechanism.

It has been reported that the W354A, E351A/D356A and E351A/W354A/D356A mutations inhibit interactions between FlhA_C_ and flagellar chaperones in complex with their cognate filament-type substrates^13^, suggesting that FlhA_L_ regulates the binding of flagellar chaperones to FlhA_C_. The crystal structure of FlhA_C_(E351A/D356A) showed that Trp-354 of one Mol-A molecule bound to the hydrophobic dimple of the flagellar chaperone binding site of its nearest Mol-A in the crystal (Fig. 4). Although the relative orientations of these Mol-A molecules in the crystal differs from those in the FlhA_C_ nonameric ring, FlhA_L_ can bind to the hydrophobic dimple of FlhA_C_ in the nonamer ring structure because of a highly flexible nature of FlhA_L_ (Fig. 5). The C-terminal region of FlhA_L_ is flexible enough to allow such subunit orientations without changing the essential interaction between FlhA_L_ and the chaperone binding site of FlhA_C_, as it has been shown to have various conformations in the known FlhA_C_ structures^18^. Therefore, we propose that an interaction between FlhA_L_ and the hydrophobic dimple of its neighboring FlhA_C_ subunit suppresses the docking of flagellar chaperones to the FlhA_C_ ring platform during HBB assembly and that the hook completion induces the detachment of FlhA_L_ from the dimple through an interaction between FliK_C_ and FlhB_C_ and its attachment to the D1 and D3 domains to induce structural remodeling of the FlhA_C_ ring, thereby terminating hook assembly and initiating filament formation (Fig. 5). Because Trp-354 of FlhA_L_ stabilized a more compact conformation of the FlhA_C_(E351A/D356A) monomer compared to the wild-type FlhA_C_ and FlhA_C_(W354A) monomers (Fig. 3), it is also possible that FlhA_L_ may block the docking of the flagellar chaperones to FlhA_C_ by covering the binding site of the same FlhA_C_ molecule.

**Fig. 5.**
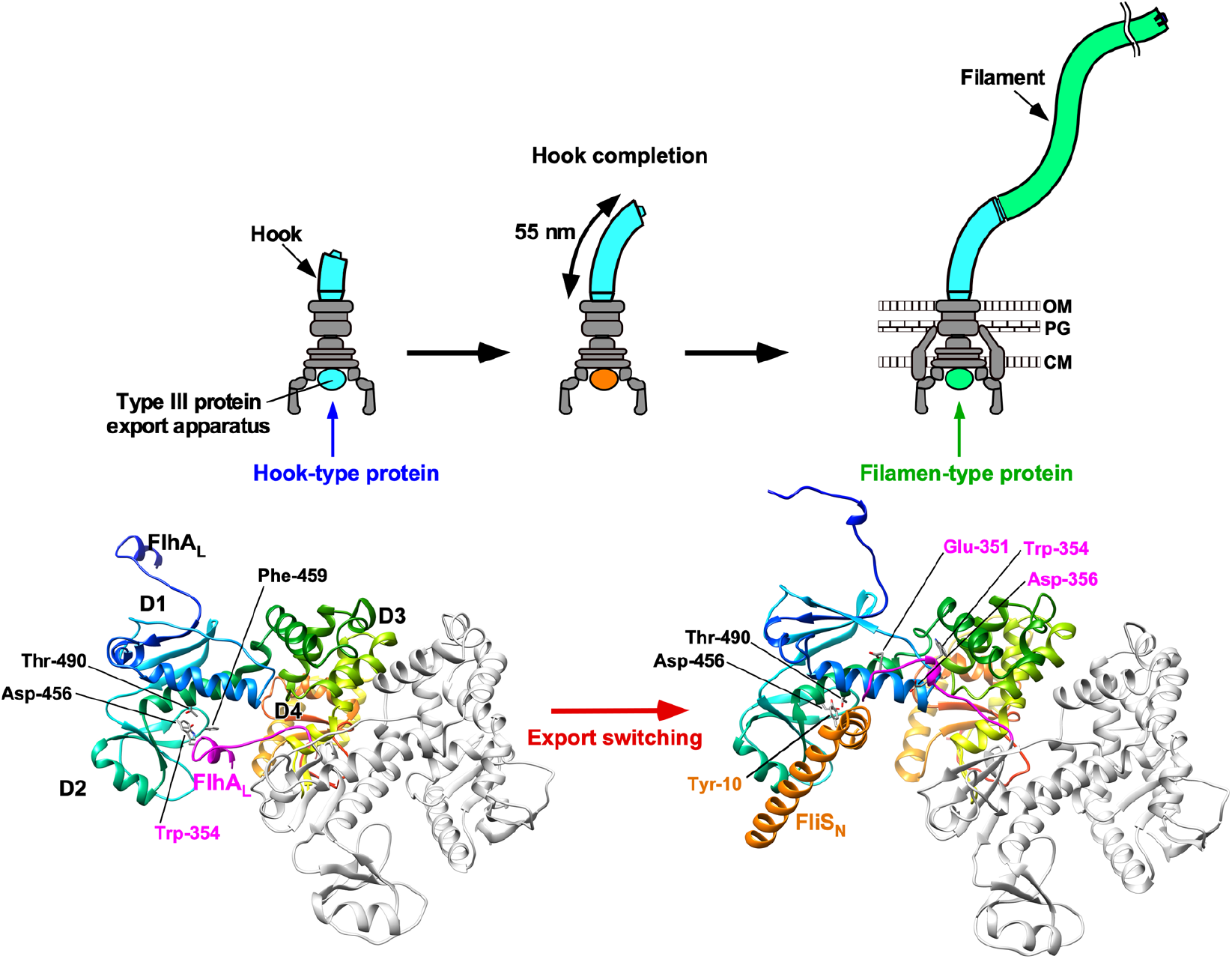
Structural rearrangements of FlhA_L_ responsible for export switching of fT3SS. Trp-359 of FlhA_L_ binds to a well-conserved hydrophobic dimple containing Asp-456, Phe-459 and Thr-490 of its neighboring FlhA_C_ subunit in the FlhA_C_ ring to inhibit the interaction of FlhA_C_ with flagellar chaperones in complex with their cognate filament-type substrates during hook assembly. When the hook reaches its mature length of about 55 nm, an interaction between FliK_C_ and FlhB_C_ triggers a conformational rearrangement of the FlhA_C_ ring so that FlhA_L_ dissociates from the hydrophobic dimple and binds to the D1 and D3 domains of the neighboring FlhA_C_ subunit, allowing the chaperones to bind to FlhA_C_ to facilitate the export of their cognate substrates for filament assembly.

## Methods

### Bacterial strains and plasmids

Bacterial strains and plasmids used in this study are listed in Supplementary Table 1.

### DNA manipulations

DNA manipulations and site-directed mutagenesis were carried out as described previously^24^. DNA sequencing reactions were carried out using BigDye v3.1 (Applied Biosystems) and then the reaction mixtures were analyzed by a 3130 Genetic Analyzer (Applied Biosystems).

### Motility assays

Fresh colonies were inoculated into soft agar plates [1% (w/v) triptone, 0.5% (w/v) NaCl, 0.35% Bacto agar] and incubated at 30°C.

### Secretion assays

Details of sample preparations have been described previously^25^. After SDS-polyacrylamide gel electrophoresis (PAGE), immunoblotting with polyclonal anti-FlgD, anti-FlgE, anti-FlgK, or anti-FliD antibody was carried out as described previously^26^.

### Hook length measurements

The HBBs were purified from NH004gM carrying pMM130 or pYI003 as described previously^27^. The HBBs were negatively stained with 2%(w/v) uranyl acetate. Electron micrographs were recorded with a JEM-1011 transmission electron microscope (JEOL, Tokyo, Japan) operated at 100 kV and equipped with a F415 CCD camera (TVIPS, Gauting, Germany). Hook length was measured by ImageJ version 1.48 (National Institutes of Health).

### Protein purification

*E. coli* BL21 (DE) Star cells carrying a pET15b-based plasmid encoding His-FlhA_C_ or its mutant variants were grown overnight at 30°C in 250 mL of L-broth [1% (w/v) tryptone, 0.5% (w/v) yeast extract, 0.5% (w/v) NaCl] containing ampicillin. His-FlhA_C_ and its mutant variants were purified by affinity chromatography, followed by size exclusion chromatography as described previously^24^.

### Far-UV CD spectroscopy

Far-UV CD spectroscopy of His-FlhA_C_ or its mutant variants was carried out at room temperature using a Jasco-720 spectropolarimeter (JASCO International Co., Tokyo, Japan) as described previously^28^.

### Cystein modification by mPEG-maleimide

His-FlhA_C_(F459C), His-FlhA_C_(K548C), His-FlhA_C_(F459C/K548C), His-FlhA_C_(W354A/F459C/K548C), His-FlhA_C_(E351A/D356A/F459C/K548C) and His-FlhA_C_(E351A/W354A/D356A/F459C/K548C) were dialyzed overnight against PBS (8 g of NaCl, 0.2 g of KCl, 3.63 g of Na_2_HPO_4_ 12H_2_O, 0.24 g of KH_2_PO_4_, pH 7.4 per liter) at 4°C, followed by cysteine modification by mPEG-maleimide (Fluka) as described previously^18^. Each protein solution was run on SDS-PAGE and then analyzed by Coomassie Brilliant blue (CBB) staining.

### X-ray crystallographic study of FlhA_C_(E351A/D356A)

Initial crystallization screening was performed at 20°C by the sitting-drop vapor-diffusion method using Wizard Classic I and II, Wizard Cryo I and II (Rigaku Reagents, Inc.), Crystal Screen and Crystal Screen 2 (Hampton Research). Crystals suitable for X-ray analysis were obtained from drops prepared by mixing 0.5 μL protein solution with 0.5 μL reservoir solution containing 0.1 M Tris-HCl (pH 8.5), 20% (v/v) PEG 8000, and 200 mM MgCl_2_. X-ray diffraction data were collected at synchrotron beamline BL41XU in SPring-8 (Harima, Japan) with the approval of the Japan Synchrotron Radiation Research Institute (JASRI) (Proposal No. 2016B2544 and 2018A2568). The FlhA_C_(E351A/D356A) crystal was soaked in a solution containing 90% (v/v) of the reservoir solution and 10% (v/v) glycerol for a few seconds and was directly transferred into liquid nitrogen for freezing. The X-ray diffraction data were collected under nitrogen gas flow at 100 K. The diffraction data were processed with MOSFLM^29^ and were scaled with Aimless^30^. The initial phase was determined by molecular replacement using the software package Phenix^31^ with the wild-type FlhA_C_ structure in the orthorhombic crystal form (PDB code: 6AI0) as a search model. The atomic model was constructed with COOT^32^ and refined with Phenix^31^. During the refinement process, iterative manual modification was performed. The diffraction data statistics and refinement statistics are summarized in Table 1.

## Accession code

The atomic coordinates have been deposited in Protein Data Bank under the accession code 7CTN.

## Acknowledgements

We thank beamline staffs at SPring-8 for technical help in use of beamlines BL41XU. This work was supported in part by JSPS KAKENHI Grant Numbers JP18K14638 and JP20K15749 (to M.K.), JP16J01859 (to N.T.), 25000013 (to K.N.), 15H02386 (to K.I.) and JP26293097 and JP19H03182 (to T.M.). This work has also been partially supported by JEOL YOKOGUSHI Research Alliance Laboratories of Osaka University to K.N.

## Author Contributions

K.N. K.I. and T.M. conceived and designed research; Y.I., M.Kida, M.Kinoshita, N.T., K.I. and T.M. preformed research; Y.I., M.Kida, M.Kinoshita, N.T., K.I. and T.M. analysed the data; and K.N., K.I. and T.M. wrote the paper based on discussion with other authors.

## Data availability

All data generated during this study are included in this published article and its Supplementary Information files.

