## Supplementary Figures 1-3 and Table 1 for "The FlhA linker mediates flagellar protein export switching during flagellar assembly"

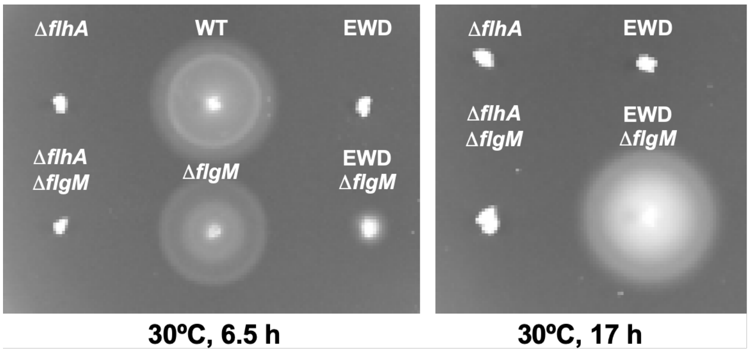

**Supplementary Fig. 1.** Motility of the *Salmonella* NH001 strain transformed with pTrc99A (Δ*flhA*), pMM130 (WT), or pYI003 (EWD) and the *Salmonella* NH001gM strain carrying pTrc99A (Δ*flhA* Δ*flgM*), pMM130 (Δ*flgM*), or pYI003 (EDW Δ*flgM*) in soft agar.

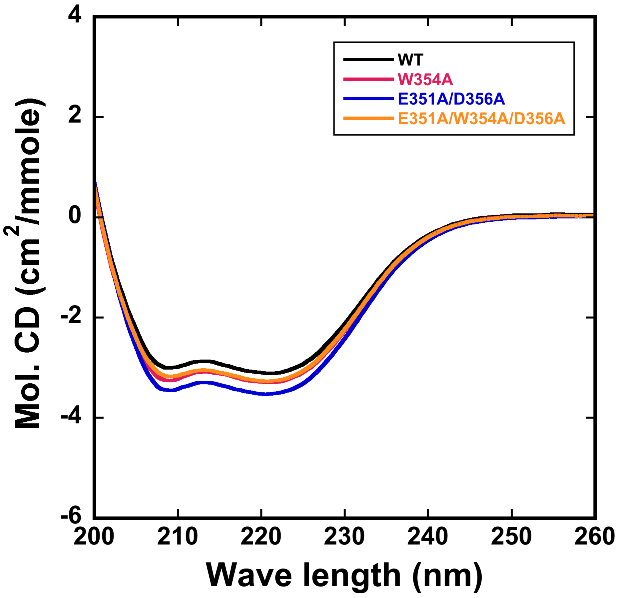

**Supplementary Fig. 2**. Effect of the W354A, E351A/D356A and E351A/W354A/D356A mutations on far-UV CD spectra of FlhA$_C$. Measurements were carried out at room temperature in 20 mM Tris-HCl, pH 8.0, in a quartz cell with a path length of 1 mm.

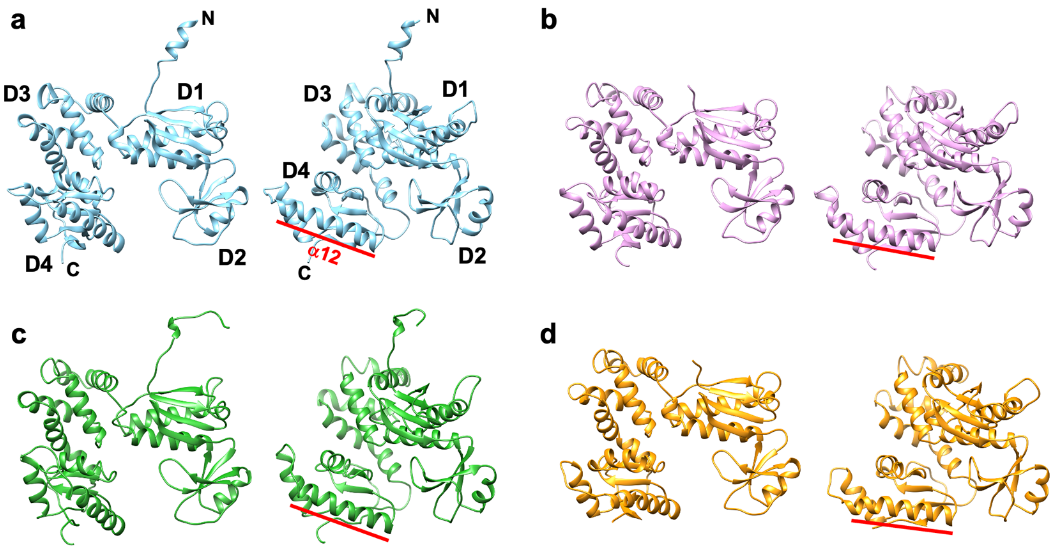

**Supplementary Fig. 3**. Two distinct conformations of FlhA$_C$(E351A/D356A) in the crystal. Ribbon representations of the crystal structures of Mol-A of FlhA$_C$(E351A/D356A) (a), Mol-B of FlhA$_C$(E351A/D356A) (b), wild-type FlhA$_C$ in an open conformation (PDB code: 3A5I) (c), and wild-type FlhA$_C$ in a semi-closed conformation (PDB code: 6AI0) (d). The right panel shows the view from the right side of the left panel. Red bar shows the direction of the α12 helix, which is a good indicator of the orientation of domain D4.

### Supplementary Table 1. Strains and plasmids used in this study.

| Strain/Plasmid | Relevant characteristics | References |
|---|---|---|
| *E. coli* | | |
| BL21 Star (DE3) | Overexpression of proteins | Novagen |
| *Salmonella* | | |
| NH001 | *ΔflhA* | [1] |
| NH001gM | *ΔflhA ΔflgM*::*km* | This study |
| YI1004-xx | Pseudorevertants isolated from NH001 carrying pYI003 | This study |
| Plasmids | | |
| pTrc99FF4 | Expression vector | [2] |
| pMM130 | pTrc99AFF4/ FlhA | [3] |
| pYI003 | pTrc99AFF4 / FlhA(E351A/W354A/D356A) | 18 |
| pYI008 | pET15b/ His-FlhA$_C$ | 13 |
| pYI009 | pET15b / His-FlhA$_C$(W354A) | 13 |
| pYI010 | pET15b / His-FlhA$_C$(E351A/D356A) | 13 |
| pYI012 | pET15b / His-FlhA$_C$(E351A/W354A/D356A) | 13 |
| pYI008(F459C) | pET15b/ His-FlhA$_C$(F459C) | 18 |
| pYI008(K548C) | pET15b/ His-FlhA$_C$(K548C) | 18 |
| pYI008(F459C/K548C) | pET15b/ His-FlhA$_C$(F459C/K548C) | 18 |
| pYI009(F459C/K548C) | pET15b/ His-FlhA$_C$(W354A/F459C/K548C) | This study |
| pYI010(F459C/K548C) | pET15b / His-FlhA$_C$ FlhA$_C$(E351A/D356A /F459C/K548C) | This study |
| pYI012(F459C/K548C) | pET15b/ His-FlhA$_C$(E351A/ W354A/D356A /F459C/K548C) | This study |
